# Placental Cell Conditioned Media Modifies Hematopoietic Stem Cell Transcriptome In Vitro

**DOI:** 10.1101/2023.03.27.534393

**Authors:** Sean M. Harris, Anthony L. Su, John F. Dou, Rita Loch-Caruso, Elana R. Elkin, Sammy Jaber, Kelly M. Bakulski

## Abstract

**Background:** Hematopoietic stem cells are cells that differentiate into all blood cell types. Although the placenta secretes hormones, proteins and other factors important for maternal and fetal health, cross-talk between placental cells and hematopoietic stem cells is poorly understood. Moreover, toxicant impacts on placental-hematopoietic stem cell communication is understudied. The goals of this study were to determine if factors secreted from placental cells alter transcriptomic responses in hematopoietic stem cells and if monoethylhexyl phthalate (MEHP), the bioactive metabolite of the pollutant diethylhexyl phthalate, modifies these effects.

**Methods:** We used K-562 and BeWo cells as *in vitro* models of hematopoietic stem cells and placental syncytiotrophoblasts, respectively. We treated K-562 cells with medium conditioned by incubation with BeWo cells, medium conditioned with BeWo cells treated with 10 μM MEHP for 24 hours, or controls treated with unconditioned medium. We extracted K-562 cell RNA, performed RNA sequencing, then conducted differential gene expression and pathway analysis by treatment group.

**Results:** Relative to controls, K-562 cells treated with BeWo cell conditioned medium differentially expressed 173 genes (FDR<0.05 and fold-change>2.0), including 2.4 fold upregulatation of *TPM4* and 3.3 fold upregulatation of *S1PR3*. Upregulated genes were enriched for pathways including stem cell maintenance, cell proliferation and immune processes. Downregulated genes were enriched for terms involved in protein translation and transcriptional regulation. MEHP treatment differentially expressed eight genes (FDR<0.05), including genes involved in lipid metabolism (*PLIN2*, fold-change: 1.4; *CPT1A*, fold-change: 1.4).

**Conclusion:** K-562 cells, a model of hematopoietic stem cells, are responsive to media conditioned by placental cells, potentially impacting pathways like stem cell maintenance and proliferation.

## Introduction

The placenta forms the critical maternal-fetal interface during pregnancy, providing the fetus with vital nutrients, gas exchange for respiration and protection from maternal immune responses (Farah et al. 2020; Kanellopoulos-Langevin et al. 2003). Throughout pregnancy, the placenta secretes a variety of factors into maternal and fetal circulation, which multiple roles in cell-cell communication via paracrine and autocrine signaling pathways (Iliodromiti et al. 2012). Cross-talk between the placenta and decidua and between placental trophoblast and placental endothelial cells has been demonstrated *in vitro* (Hess et al. 2007; Troja et al. 2014). Biologically active factors secreted by placental syncytiotrophoblasts may play a role in this cross-talk, as these large multi-nucleated cells form the outermost of layer of placental villi and serve as the major interface between fetal and maternal circulatory systems (Huppertz 2018). Major gaps remain in our understanding of the role that factors secreted by syncytiotrophoblasts play in placental development.

The human placenta is a hematopoietic organ (Barcena 2009, 2011, Robin et al 2009, Serikov 2009), acting as an early niche for hematopoietic stem cells during fetal development before hematopoiesis is taken over by the fetal liver (Dzierzak and Robin 2010). Hematopoiesis is a critical process in the development of the immune system (Jagannathan-Bogdan and Zon 2013). Production of the blood throughout the life course is dependent on hematopoietic stem cell self-renewal and differentiation into the various blood cell types (Weissman 2000). Environmental or genetic factors that disrupt the process of hematopoietic stem cell migration, proliferation, differentiation, or self-renewal, could adversely affect immune function later in life (Bao et al. 2019; Laiosa and Tate 2015). Moreover, adverse pregnancy outcomes such as preeclampsia are associated with disrupted differentiation capacity in hematopoietic cells in the umbilical cord blood and in the fetal liver (Masoumi et al. 2019; Stallmach et al. 1998). The specific placental cell types that mediate hematopoietic stem cell differentiation in the placental niche are poorly understood.

The neonatal development of blood cells is a complex process that occurs in multiple anatomical sites, which change over time. Early hematopoiesis begins in the yolk sac before proceeding to the placenta, fetal liver and finally ending with colonization of the bone marrow at birth (Dzierzak and Speck 2008; Mikkola and Orkin 2006). The placental role in early hematopoiesis is an important but understudied area of pregnancy biology (Dzierzak and Robin 2010; Robin et al. 2009). The hematopoietic stem cell microenvironment plays an important role in the maintenance of stem cell properties as shown by the loss of self-renewal capacity in hematopoietic stem cells *in vitro* (Rhodes et al. 2008). Environmental toxicants can interfere with key processes, such as hematopoietic stem cell differentiation into lymphocytes (Ahrenhoerster et al. 2014), dysregulation of differentiation pathways (Votavova et al. 2011) and erythropoiesis (Demur et al. 2013).

Toxicology researchers have also discovered that multiple environmental chemicals, such as phthalates (Tetz et al. 2013), trichloroethylene (Elkin et al. 2018), and polycyclic aromatic hydrocarbons (Drwal et al. 2020), disrupt placental cells via mechanisms such activation of apoptosis, inflammation and endocrine disruption. Diethylhexyl phthalate, which is metabolized to monoethylhexyl phthalate (MEHP) is a widespread contaminant and known endocrine disruptor with toxicologic effects in the placenta (Den Braver-Sewradj et al. 2020; Mattiske and Pask 2021; Zhang et al. 2021). Placental effects of exposure to DEHP or other phthalates include decreased placental weight in animal models (Zhang et al. 2016; Zong et al. 2015) and decreased methylation and transcription of growth-related genes in human studies (Grindler et al. 2018; Zhao et al. 2016). Another mechanism of toxicity during pregnancy is via impacts on maternal and fetal blood cells, including hematopoietic stem cells. In a 2015 review Laiosa, et al. highlighted fetal hematopoietic stem cells as a potential target of endocrine disrupting chemicals, tobacco smoke and pesticides (Laiosa and Tate 2015), noting that animal studies showed maternal exposures to chemicals such as tetrachlorodibenzo-p-dioxin (Fine et al. 1990) and nicotine suppress hematopoietic activity in the fetal liver and bone marrow (Serobyan et al. 2005). Similar effects were observed for the effects of tobacco smoke during pregnancy in women. For example, a transcriptomic study of cord blood showed that pathways involved in hematopoiesis and immune cell differentiation were downregulated in the blood cells of mothers who smoked during pregnancy (Votavova et al. 2011).

Despite playing a clear role early in development of the immune and blood systems, and despite opportunity for exposure of pregnant women to a wide range of environmental chemical exposures, the extent to which environmental toxicants disrupt interactions between placental cells and hematopoietic stem cells has scarcely been explored. The objectives of this study were two-fold: (1) to determine if factors secreted from syncytiotrophoblasts play a role in communication with hematopoietic stem cells and (2) to determine if this cell-cell communication is altered by treatment with the relevant metabolite of a widespread environmental toxicant and known endocrine disruptor, diethylhexyl phthalate (DEHP) (Den Braver-Sewradj et al. 2020; Mattiske and Pask 2021; Zhang et al. 2016).

## Methods

### Chemicals and Reagents

Iscove’s Modified Dulbecco’s Medium (IMDM), F12-K Nutrient Mixture Kaighn’s Modification with (+) L Glutamine, Dulbecco’s Modified Eagle Medium (DMEM)/F12 Nutrient Mixture without phenol red, penicillin/streptomycin (P/S), heat-inactivated fetal bovine serum (HI-FBS) and exosome-depleted fetal bovine serum (ED-FBS), which contains over 90% of exosomes depleted, were purchased from Gibco (Grand Island, NY). Phosphate buffered saline (PBS) and 0.25% trypsin-EDTA were from Invitrogen Life Technologies (Carlsbad, CA). Dimethyl sulfoxide (DMSO) was purchased from Tocris Bioscience (Bristol, United Kingdom). Forskolin and 2-mercaptoethanol were purchased from Sigma-Aldrich (St. Louis, MO). Mono-2-ethylhexyl phthalate (MEHP) was purchased from AccuStandard (New Haven, CT).

### Cell Culture

BeWo (ATCC CCL-98), a human placental trophoblast cell line (Pattillo and Gey 1968) and K-562 (ATCC CCL-243), a hematopoietic stem cell line (Lozzio and Lozzio 1975), were purchased from American Type Culture Collection (ATCC, Manassas, VA). Cells used in experiments were within twenty passage numbers from arrival into the laboratory and were routinely verified by their short tandem repeat profiles using fragment analysis (ABI 3730XL DNA Analyzer, Applied Biosystems, Waltham, MA) at the University of Michigan Advanced Genomics Core. Unless otherwise noted, all media were supplemented with 10% (v/v) HI-FBS and 1% (v/v) of 10,000 U/mL P/S. IMDM and F12-K Nutrient Mixture Kaighn’s Modification with (+) L Glutamine were used as the media for culturing for K-562 cells and BeWo cells, respectively. All cells were plated at a 100,000 cells/mL in 25 mL of media in 175 cm^2^ flasks (Corning Inc., Corning, NY) and subcultured at 70-80% confluence. Unlike BeWo cells, K-562 cells are suspension cells and do not require 0.25% trypsin-EDTA for detachment in subculture. Cell cultures were maintained in a 5% CO_2_, 37°C controlled and humidified incubator. This work with human cell cultures was approved by the University of Michigan Institutional Biosafety Committee (IBCA00000100).

### Generation of BeWo Conditioned and Unconditioned Media

Conditioned media were prepared from cultures of syncytialized BeWo cells that were treated with and without MEHP. BeWo cells were seeded into 6-well plates at 200,000 cells/well. After 24 hours, BeWo cells were treated with 100 μM forskolin for 48 hours to stimulate syncytialization (i.e., cell fusion) (Wice *et al*., 1990; Inadera *et al*., 2010). After syncytialization, BeWo cells were washed 3 times with PBS and then treated for an additional 48 hours with MEHP (10μM) or vehicle control (0.05% DMSO) in medium containing 10% (v/v) exosome depleted fetal bovine serum (ThermoFisher, Waltham, MA) and 1% (v/v) of 10,000 U/mL P/S. Exosome depleted medium was used to minimize the influence of extracellular vesicles such as exosomes, which are found in standard FBS (Kornilov et al. 2018). After treatment with MEHP or vehicle control, treatments were completed, medium was collected and stored at −80°C. Unconditioned media was prepared in the same manner as vehicle control media but had no contact with cells.

### Treatment of K-562 Cells With BeWo Conditioned Media

The experimental design for K-562 treatment with BeWo conditioned media is shown in Figure 1. K-562 cells were plated at 100,000 cells/mL in Corning 6-well plates. After 72 hours in culture, cells were washed 3 times with PBS and treated with either: 1) unconditioned media, 2) BeWo conditioned media or 3) MEHP+BeWo conditioned media. There were four replicates for each of the three groups.

**Figure 1.**
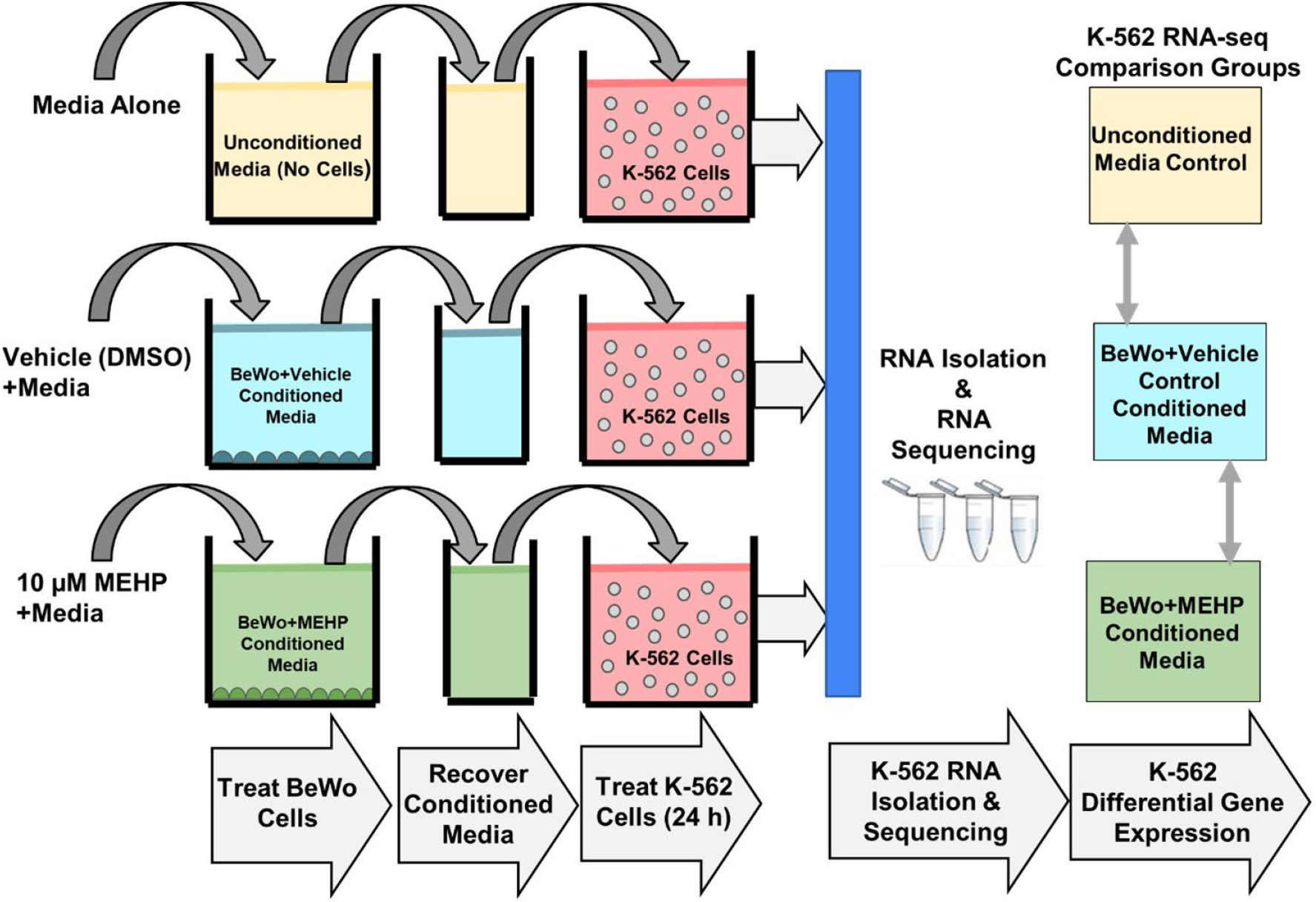
Experimental design. K-562 cells were treated with unconditioned medium, BeWo conditioned medium or MEHP (10μM) + BeWo conditioned medium. After 24 hours of treatment, RNA was isolated from K-562 cells followed by RNA sequencing and differential gene expression analysis.

### RNA Extraction

RNA was extracted from K-562 cells across all three treatment groups (unconditioned media, BeWo conditioned media and MEHP (10μM) + BeWo conditioned media) using the RNeasy PLUS Mini Kit from Qiagen (Germantown, MD). RNA extraction followed the manufacturer instructions, with the addition of a 2-minute spin at 15,000 x g in the QIAshredder (Qiagen, Germantown, MD) prior to the genomic DNA elimination step. Buffer RLT Plus was supplemented with 1% (v/v) 2-mercaptoethanol to ensure that solutions were free of RNases. RNA purity and concentration were determined using the NanoDrop 2000 UV-Vis Spectrophotometer (Thermo Fisher Scientific, Waltham, MA). RNA was stored at −80°C until further analysis.

### Sequencing

RNA sequencing was performed at the University of Michigan Advance Genomics Core. Stranded sequencing libraries for RNA isolated from K-562 cells were prepared with the TruSeq Stranded mRNA Library Prep Kit (lllumina, San Diego, CA). Libraries were sequenced on one lane using single-end 50 cycle reads on a HiSeq 4000 sequencer (lllumina).

### RNA-seq Processing

Raw fastq files were first examined using fastQC (version 0.11.5) (Andrews 2010), and reports generated for the 12 samples were collated using multiQC (version 0.9) (Ewels et al. 2016). All samples had sequences of 51 base pairs in length. Mean quality scores across all base positions were high across samples. Per base sequence content was not balanced for the first bases. For example, across all samples there was ~60% G at the first base, and ~35% T at the second. GC content was 48% or 49% for all samples. Overrepresented sequences made up less than 1% of all reads in all 12 samples, and adapter contamination >0.1% was not found. However, samples had between 63% and 75% of reads duplicated. Approximately 20% to 25% of sequences had a sequence duplication level between 10 and 50, while 13% to 24% of reads were not duplicated. We mapped reads to the human reference genome (hg38) using the Spliced Transcripts Alignment to a Reference (STAR) (version 2.6.0c) program (Dobin et al. 2013). Post alignment, we used QoRTs (version 1.3.6) (Hartley and Mullikin 2015) to examine further quality control metrics. Sample distributions in quality control metrics were similar to each other, with no extreme outliers. Next, featureCounts (version 1.6.1) (Liao et al. 2014) quantified these aligned reads. We used default behavior to drop multi-mapping reads and count features mapping to exons.

### Differential Gene Expression Analysis

Following alignment and quantification, we tested for differentially expressed genes. Gene counts were read into R (version 3.6.0), which we analyzed with the DESeq2 package (version 1.24.0) (Love et al. 2014). We plotted principal components, calculated on variance stabilizing transformed values of the expression data, to examine clustering. Plots were painted by treatment group and by laboratory day to assess potential batch effects. In DESeq2, our model terms were treatment group (three levels: 1) unconditioned media, 2) BeWo conditioned media or 3) MEHP (10μM) + BeWo conditioned media) and day of sample culture. We used an adjusted p-value (false discovery rate, FDR) < 0.05 and an absolute log_2_(fold-change) > 1.0 to determine significance.

We examined two contrasts of interest. To investigate the effect of the placental media, we compared the BeWo conditioned media to unconditioned media. To investigate the effect of phthalate, we compared the MEHP (10μM) + BeWo conditioned media group to the BeWo conditioned media group. Default settings for DESeq2 were used for filtering of genes with low normalized mean counts. We created volcano plots of results using the EnhancedVolcano (version 1.2.0) package, after applying log fold change shrinkage using the “apeglm” prior (Zhu et al. 2019).

### Pathway Analysis

We used RNA-enrich to identify significantly enriched gene sets among genes changed in cells treated with BeWo conditioned media (Lee et al. 2016). RNA-enrich tests for enrichment for relevant biological concepts across several databases, including gene ontology biologic processes terms, Medical Subject Headings (MeSH), Kyoto Encyclopedia of Genes and Genomes (KEGG) pathways, Drug Bank, and Metabolite annotations. We used the directional method in RNA-enich, which allows for the discrimination between biological terms enriched with either upregulated or downregulated genes. We identified significantly altered biological concepts using a cutoff of FDR<0.05 and odds ratio >1.1 or <0.9. We then used REVIGO (Supek et al. 2011) to remove redundant gene ontology terms for the list of significantly enriched terms, using the default REVIGO settings except gene ontology list size was set to “Small”. We did not conduct pathway enrichment analysis for MEHP (10μM) + BeWo conditioned samples due to the low number of significant gene expression changes in this treatment group.

### Data and Code Availability

Code to complete all analyses is publicly available (www.github.com/bakulskilab). RNA expression data are publicly available through the genome expression omnibus (accession # GSE188187).

## Results

### Sample Sequencing Descriptive Statistics

Our experiment consisted of three treatments: 1) unconditioned media, 2) BeWo-conditioned medium or 3) MEHP + BeWo-conditioned medium. Each group had four samples. Following alignment and quantification, samples had between 18,333,598 to 32,618,806 reads assigned to features numbering from 20,689 to 22,182 (Supplementary Table 1). In principal component plots, we observed clustering by treatment group, and by culture date (Supplementary Figure 1).

### BeWo conditioned media: Differential gene expression in K562 cells

We evaluated the effect of BeWo conditioned medium by examining genes differentially expressed between the BeWo-conditioned group and the unconditioned medium group. Following filtering of genes with low normalized mean counts, 14851 genes were analyzed. Treatment with BeWo-conditioned medium differentially expressed 3743 genes using statistical criteria (FDR<0.05), 174 genes by fold change criteria (log_2_(fold change) > 1.0), and 173 genes met both criteria (Figure 2). Of genes meeting both criteria, 115 (66%) were upregulated with BeWo conditioned media treatment. BeWo conditioned media treatment upregulated the following, with the smallest adjusted p-values: Tropomyosin 4 (*TPM4*, fold-change: 2.4, adjusted-p=1.8×10^-53^), Sphingosine-1-Phosphate Receptor 3 (*S1PR3*, fold-change: 3.3, adjusted-p=1.6×10^-40^), Jun Proto-Oncogene, AP-1 Transcription Factor Subunit (*JUN*, fold-change: 2.4, adjusted-p=5.3×10^-28^) and Ring Finger Protein 144A (*RNF144A*, fold-change: 2.0, adjusted-p=7.3×10^-26^). BeWo conditioned media treatment upregulated the following, with the largest fold-change values: Glycoprotein Hormones, Alpha Polypeptide (*CGA*, fold-change: 1,552, p-adjusted=4.5×10^-19^), Chorionic Gonadotropin Subunit Beta 8 (*CGB8*, fold-change: 91, adjusted-p=1.1×10^-4^), Collagen type IV alpha 1 chain (*COL4A1*, fold-change: 91, adjusted-p=8.6×10^-6^) and Aquaporin 6 (*AQP6*, fold-change: 79, adjusted-p=1.2×10^-4^). The full list of genes and differential expression results are shown in Supplementary Table 2.

**Figure 2.**
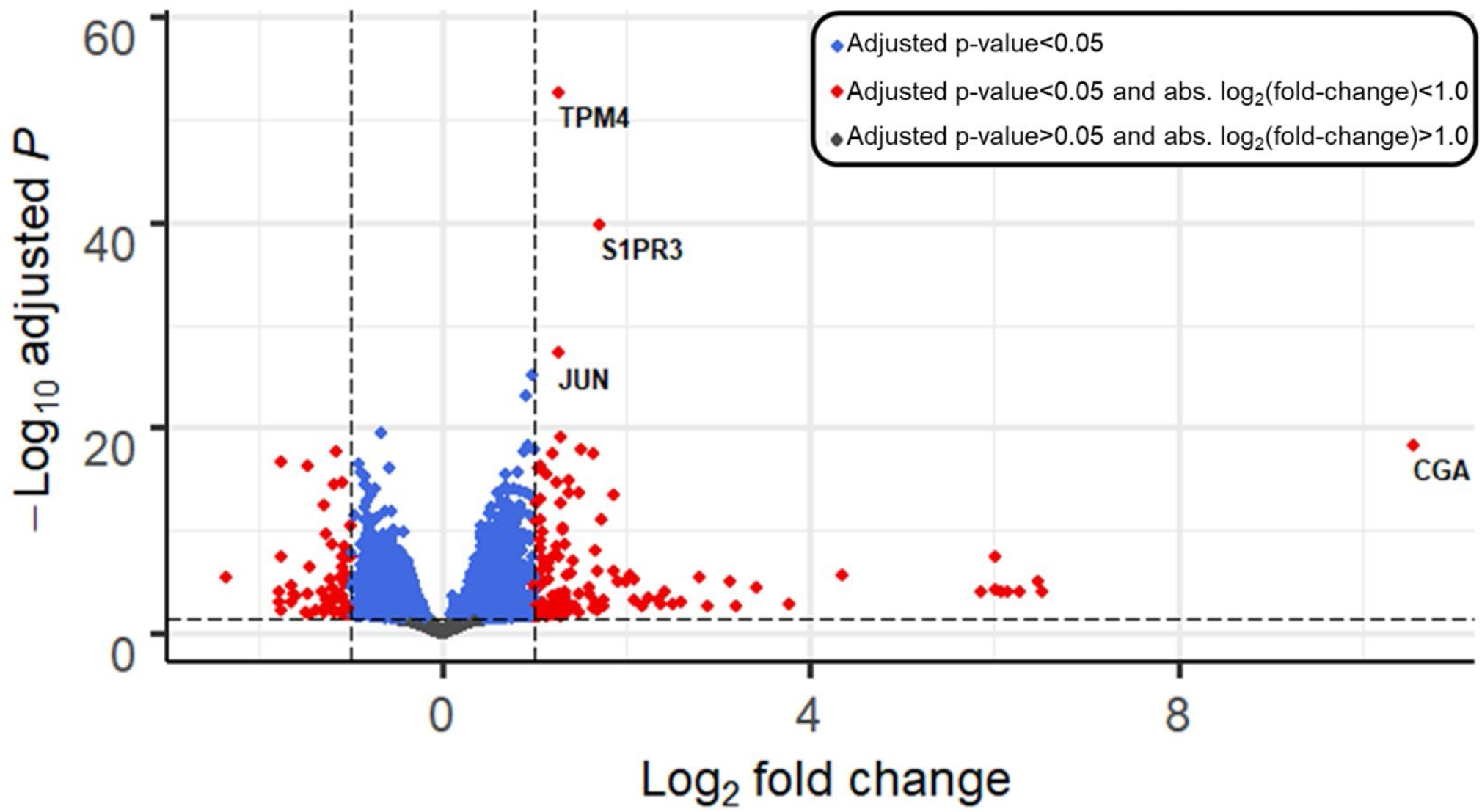
Volcano plot depicting differential gene expression in K562 cells after 24-h treatment with BeWo-conditioned medium vs. compared to unconditioned medium (control). Genes are plotted by log_2_(fold change) (x-axis), and –log_10_(adjusted p-values) (y-axis). Horizontal line shows significance cutoff (adjusted p-value<0.05). Vertical lines show fold-change cutoffs of absolute log_2_(fold-change) > 1. Blue points have adjusted p-values < 0.05. Red points adjusted p-values < 0.05 plus log-fold change > 1.0.

### Pathway Analysis of Genes Impacted by BeWo-conditioned Medium

After removing redundant gene ontology terms, we identified 70 pathways that were significantly (FDR < 0.05) enriched among upregulated genes and 63 pathways enriched among down regulated genes in K-562 cells treated with BeWo-conditioned medium compared to unconditioned medium. Upregulated pathways were involved with biological functions including stem cell maintenance (“somatic stem cell population maintenance”, FDR=0.001), cell migration (“positive regulation of cell migration”, FDR=9.4*10^-7^), immune or inflammatory processes (“cytokine secretion”, FDR=0.001), tissue/organ system development (“regulation of vasculature development”, FDR=0.0002), cell signaling pathways (“phosphatidylinositol 3-kinase signaling”, FDR=0.007) and embryonic development (“formation of primary germ layer”, FDR=1.4*10^-5^) (Figure 3). Downregulated pathways were involved with biological functions including protein translation (“mitochondrial translation”, FDR=0.0001; “translational elongation”, FDR=0.002), transcriptional processes (“RNA processing”, FDR=3.7*10^-14^), immune or inflammation processes (“regulation of interleukin-6 biosynthetic process”, FDR=0.01) and metabolism (“gluconeogenesis”, FDR=0.04). The full list of pathway enrichment results is shown in Supplementary Table 3.

**Figure 3.**
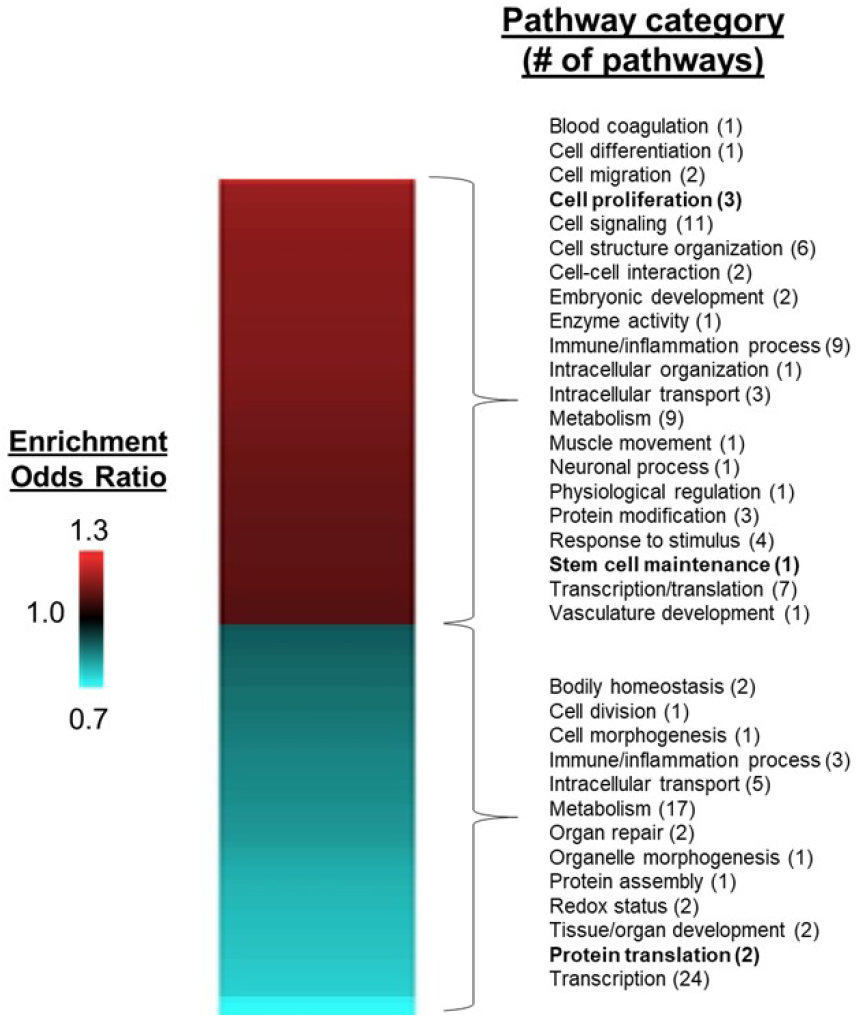
Heatmap of significantly enriched pathways for K-562 cells treated with BeWo conditioned media, compared with unconditioned medium controls. After 24-hour incubation with conditioned media, RNA was isolated from K-562 cells and used for RNA sequencing, followed by differential gene and pathway enrichment analysis. Pathways significantly enriched with upregulated genes (FDR < 0.05, enrichment odds ratio > 1.1) are shown in red and pathways enrich with downregulated genes are shown in blue (FDR < 0.05, enrichment odds ratio < 0.9). Pathway categories of particular relevance to stem cell biology are shown in bold.

### Phthalate Treatment: Differential Gene Expression

We evaluated whether the phthalate MEHP modified the effect of BeWo-conditioned medium by examining differential gene expression, comparing the MEHP + BeWo-conditioned group and the BeWo-conditioned medium group. After filtering for low normalized mean counts a total of 30760 genes remained in analysis. There were 8 genes with adjusted p-value < 0.05, and 5 genes with log-fold change > 1.0. No genes met both of these conditions (Figure 4). Genes with adjusted p-values < 0.05 included: Perilipin 2 (*PLIN2*, fold-change: 1.4, adjusted-p=3.2×10^-5^), Small Proline Rich Protein 2B (*SPRR2B*, fold-change: 0.99, adjusted-p=0.003), Transferrin Receptor (*TFRC*, fold-change: 1.3, adjusted-p=0.003), Lnc-MASTL-3 (*ENSG00000262412*, fold-change: 0.99, adjusted-p=0.003), Calponin 2 Pseudogene 1 (*CNN2P1*, fold-change: 0.99, adjusted-p=0.005), Dehydrogenase/Reductase 2 (*DHRS2*, fold-change: 0.8, adjusted-p=0.01), Carnitine Palmitoyltransferase 1A (*CPT1A*, fold-change: 1.4, adjusted-p=0.02) and Chloride Channel Accessory 2 (*CLCA2*, fold-change: 1.0, adjusted-p: 0.04). The full list of genes and differential expression results for MEHP treatment is shown in Supplementary Table 4. We observed too few differentially expressed genes to perform pathway enrichment analyses.

**Figure 4.**
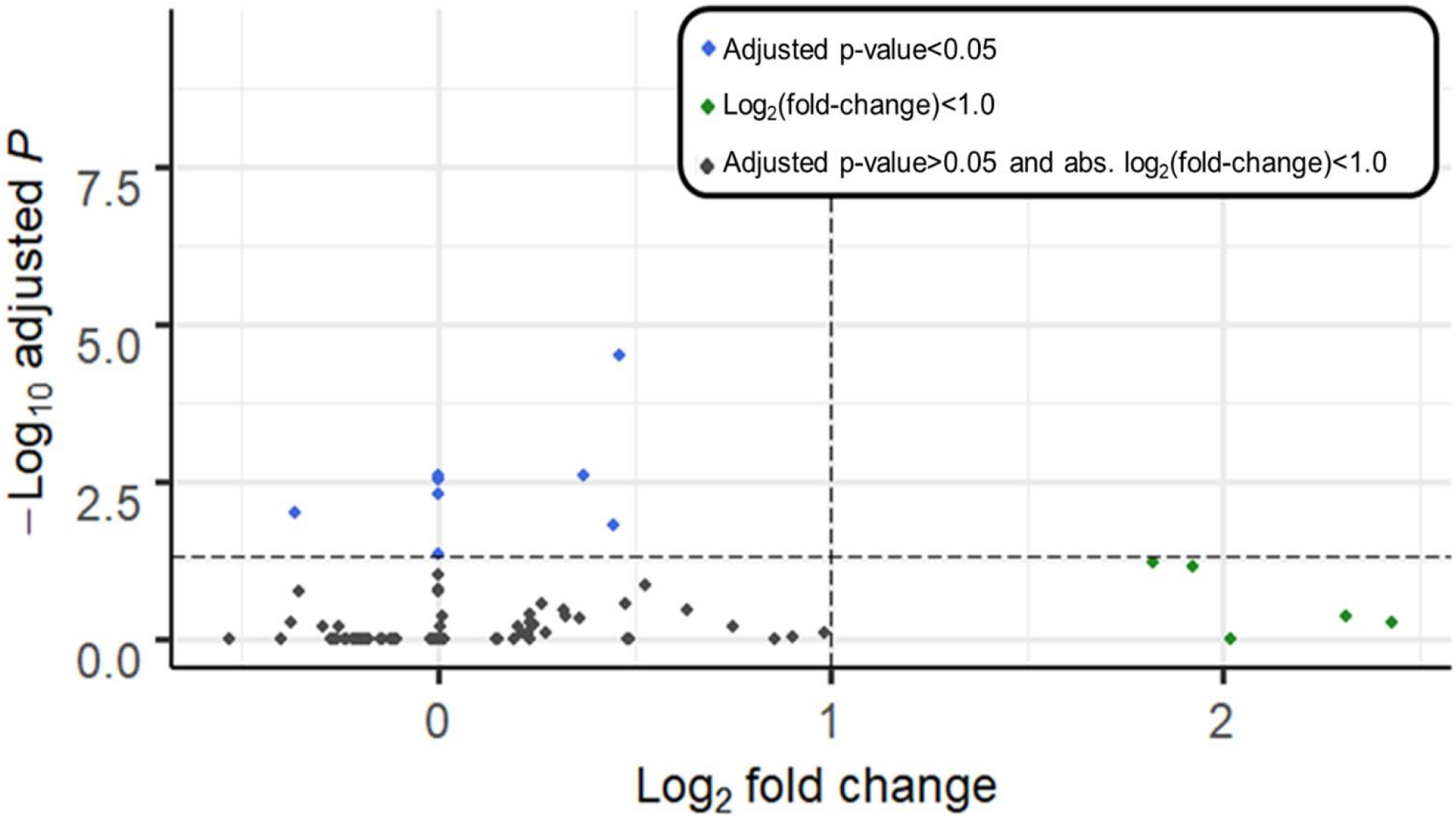
Volcano plot depicting differential gene expression of K562 cells treated with BeWo+MEHP-conditioned medium for 24 hours, compared with BeWo-conditioned medium (vehicle control with no MEHP). Genes are plotted by log_2_(fold change) (x-axis), and −log_10_(adjusted p-values) (y-axis). Horizontal line shows significance cutoff (adjusted p-value<0.05). Blue points have adjusted p-value < 0.05 Vertical lines show fold-change cutoffs (absolute log_2_(fold-change) > 1). Green points have log-fold change > 1.0.

## Discussion

This study shows that an *in vitro* model of hematopoietic stem cells (K-562) is responsive to media that has been conditioned by placental cells, potentially impacting processes related to stem cell maintenance and proliferation. These findings have important implications for communication between placental cells and hematopoietic stem cells. The placenta is an early site of fetal hematopoiesis, and placental hematopoietic stem cells are involved in the early stages of fetal blood cell differentiation. Understanding the role that specific placental cell types play in maintaining the hematopoietic stem cell environment microenvironment is critical to understanding the mechanisms underlying disorders of the immune system which may be rooted in early life events such as *in utero* exposures to environmental toxicants. Our findings suggest that synctytiotrophoblasts, the hormonally active, large multi-nucleated cells that line the outer layer of placental villi, play an important role in maintaining stem cells niche for hematopoietic stem cells during early periods of hematopoiesis in the placenta.

Previous have shown that placental hematopoietic stem cells in the mouse placenta were multipotetnial and highly proliferative, whereas hematopoietic cells in the fetal liver were unilineage suggesting that the placenta is a unique niche for hematopoietic stem cells (Gekas et al. 2005). In this study, we found that K-562 cells (a model of human hematopoietic stem cells) treated with media conditioned with differentiated BeWo cells (a model of human placental syncytiotrophoblasts) contained altered gene expression patterns compared to unconditioned controls. Pathway analysis revealed that multiple pathways were upregulated by BeWo conditioned-media including those involved in stem cell maintenance (“somatic stem cell population maintenance”) and cell proliferation (“G0 to G1 transition” and “positive regulation of endothelial cell proliferation”). Our findings suggest that syncytiotrophoblasts secrete factors that supply microenvironmental cues to hematopoietic stem cells as they move through the placental niche (Mikkola and Orkin 2006) and are consistent with the placental microenvironment maintaining hematopoietic stem cell populations in a proliferative and undifferentiated state (Gekas et al. 2005).

We also observed multiple enriched pathways with downregulated genes stimulated by treatment with BeWo conditioned media. Importantly, many of these pathways were associated with immune and inflammatory biological processes (“interleukin-6 biosynthetic process” and “regulation of interleukin-6 biosynthetic process”). Downregulation of the genes in these pathways may indicate a role for placental cells in mediating cell and/or tissue specification during fetal development. For example, inflammatory signaling via pro-inflammatory cytokine (IL-6, TNF-α) stimulation of the NF-κB and STAT3 pathways, is known to play a role in directing hematopoietic stem cell specification (King and Goodell 2011; Pietras 2017). In addition to immune and inflammatory processes, genes involved in enriched RNA translation pathways (“translational termination”, “translational elongation” and “mitochondrial translation”) were also downregulated. This further suggests a role for placental cells in maintaining hematopoietic stem cells in an undifferentiated state while in the placental niche because suppressed translation is necessary to maintain undifferentiation (Signer et al. 2014; Tahmasebi et al. 2018).

Our findings are consistent with earlier studies which showed that the placental microenvironment plays a role in directing the development of hematopoietic stem cells. For example, stromal cell lines derived from human placenta support hematopoiesis when co-cultured with human umbilical cord cells (2009). Our results suggest that other placental cell types such as syncytiotrophoblasts may also play a role in supporting hematopoiesis during fetal development. Future experiments will investigate additional placental cell types and determine which cell types are involved in the maintenance of stem cell populations in the placental microenvironment during fetal development.

We selected the a bioactive metabolite of the toxicant DEHP for this study due to widespread human exposures and known toxic effects on placental cells and placental development, including decreased placental weight in exposed mice and decreased hCGβ release from villous cytotrophoblasts (Gao et al. 2017; Martinez-Razo et al. 2021; Shoaito et al. 2019; Zhang et al. 2016). The concentration of MEHP used in this study was selected based on studies showing impacts on placental cells such as inhibition of extravillous trophoblast invasion (Gao et al. 2017) and endocrine disruption (decreased hCGβ release) (Shoaito et al. 2019) at 10 μM concentrations without impacts on cell viability. Treatment with MEHP at this concentration had a modest effect on K-562 gene expression responses, with eight statistically significant gene expression changes between treated and non-treated samples. Differentially expressed genes were involved in processes such as fat/lipid metabolism (*PLIN2*, fold-change: 1.4; *CPT1A*, fold-change: 1.4), consistent with earlier studies showing that MEHP disrupts lipid metabolism and upregulates gene targets of peroxisome proliferator-activated receptor gamma (PPARγ) (Chiang et al. 2017; Jia et al. 2016; Posnack et al. 2012; Wang et al. 2020). Future experiments should investigate MEHP effects on additional doses and time points to fully assess potential impacts on syncytiotrophoblast-hematopoietic stem cell communication.

This study had several limitations that should be noted. We used an *in vitro* cell culture study design to conduct the experiments reported here, which may not accurately reflect *in vivo* conditions because tissue structure and cell to cell interactions are lost. Both BeWo cells and K-562 cells are cell lines that originate from cancers, including choriocarcinoma for BeWo (Pattillo and Gey 1968) and chronic myelogenous leukemia for K-562 (Lozzio and Lozzio 1979; Lozzio and Lozzio 1975). Tumorigenesis inevitably changes some cellular characteristics relative to primary cells, including the ability to divide indefinitely under cell culture conditions. Despite these limitations, BeWo cells have been used extensively used to model syncytiotrophoblasts *in vitro* (Gohner et al. 2014; Hannan et al. 2010) and K-582 cells have been used to model hematopoietic stem cells (Andersson et al. 1979). Moreover, cell lines are a useful tool because of their availability, minimal time investment and low cost (Gohner et al. 2014). Future studies can test the applicability of these findings to primary cells and tissues. Finally, this study examined the effects of BeWo-conditioned media and did not isolate any of the specific biological factors known to be secreted by placental cells. These factors include hormones (Iliodromiti et al. 2012), proteins (Michelsen et al. 2019), and microvesicles (Tong and Chamley 2015), which can influence fetal development and/or maternal homeostasis during pregnancy. Future experiments could identify specific proteins, hormones or other factors that play key roles in syncytiotrophoblast-hematopoietic stem cell communication.

In conclusion, this preliminary study shows that signaling between syncytiotrophoblasts-hematopoietic stem cells could play an important role in mediating hematopoiesis while hematopoietic stem cells are in the placental niche. Importantly, understanding mechanisms that underlie the development of the immune system, specifically blood stem cells, during the sensitive fetal lifestage has important implications for adverse pregnancy outcomes or blood disorders later in life. Our findings support the role of syncytiotrophblasts in maintaining hematopoietic stem cell properties during this critical period.

## Acronyms

MEHP: monoethylhexyl phthalate
FDR: false discovery rate

## Disclosure of Interests

The authors declare no conflicts of interest.

## Funding

This work was supported by the NIH National Institute of Environmental Health Sciences (Grant Nos. P42 ES017198, P30 ES017885, and T32 ES007062), NIH National Institute of Diabetes and Digestive Kidney Diseases (Grant No. T32 DK071212) and Michigan Institute for Clinical and Health Research funded by the NIH National Center for Advancing Translational Sciences (Grant No. UL1 TR002240) Additional training grant fellowship support for ALS was from the Eunice Kennedy Shriver National Institute of Child Health and Human Development (NICHD), NIH (T32HD079342). The content is solely the responsibility of the authors and does not necessarily represent the official views of the NICHD, NIEHS, NIDDK, or NIH.

## Acknowledgments

We thank the University of Michigan Advanced Genomics Core for conducting RNA sequencing for this study.

## Notes

### Competing Interest Statement

The authors have declared no competing interest.

