## Supplemental File 1 for "Placental Cell Conditioned Media Modifies Hematopoietic Stem Cell Transcriptome In Vitro"

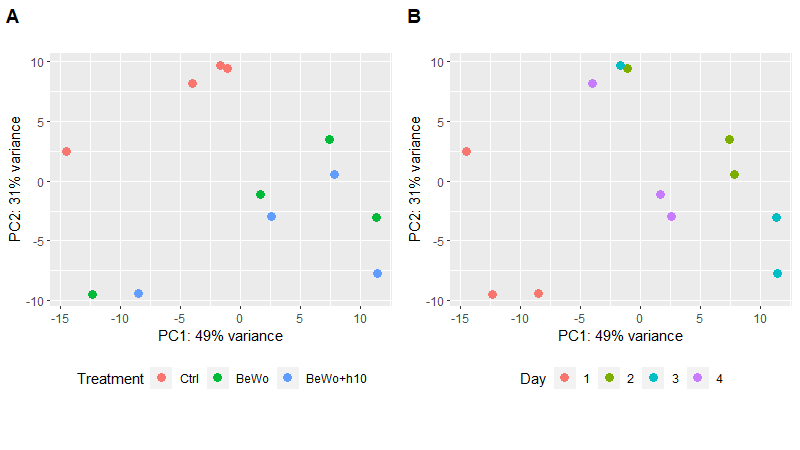
**Supplementary Figure 1**. **Plots of first two principal components calculated on variance stabilizing transformed count data.** Points of plots are colored by treatment group (A) and day of culture (B). Groups, each with four cultures, were control (Ctrl), treatment with BeWo placenta media (BeWo), and treatment with BeWo placenta media and phthalate (BeWo+h10). Cultures were done on four days, each day having one of each treatment.
