## Supplemental Table 1 for "Placental Cell Conditioned Media Modifies Hematopoietic Stem Cell Transcriptome In Vitro"

**Supplementary Table 1**. Sample sequencing statistics. Number of reads per sample did not differ by treatment group (ANOVA p-value = 0.89), nor did number of genes per sample (ANOVA p-value = 0.77).

| **Sample** | **Treatment Group** | **Day** | **N Reads** | **N Genes** |
| --- | --- | --- | --- | --- |
| 119411 | vc_minusBeWo | 1 | 25729309 | 22182 |
| 119414 | vc_minusBeWo | 2 | 22235574 | 20697 |
| 119417 | vc_minusBeWo | 3 | 28526592 | 21841 |
| 119420 | vc_minusBeWo | 4 | 24369628 | 21291 |
| 119412 | vc_plusBeWo | 1 | 24831372 | 22040 |
| 119415 | vc_plusBeWo | 2 | 25213708 | 20689 |
| 119418 | vc_plusBeWo | 3 | 23110291 | 20840 |
| 119421 | vc_plusBeWo | 4 | 22558203 | 21353 |
| 119413 | h10_plusBeWo | 1 | 24152687 | 22010 |
| 119416 | h10_plusBeWo | 2 | 32618806 | 21508 |
| 119419 | h10_plusBeWo | 3 | 23076608 | 21404 |
| 119422 | h10_plusBeWo | 4 | 18333598 | 20939 |
